# mRNA compartmentalisation spatially orients tissue morphogenesis

**DOI:** 10.1101/374850

**Authors:** Guilherme Costa, Joshua Bradbury, Nawseen Tarannum, Shane P. Herbert

**Affiliations:** Faculty of Biology, Medicine and Health, Michael Smith Building, University of Manchester, Oxford Road, Manchester, M13 9PT, UK

## Abstract

Polarised targeting of diverse mRNAs to motile cellular protrusions is a hallmark of cell migration^1–3^. Although a widespread phenomenon, definitive functions for endogenous targeted mRNAs and their relevance to modulation of *in-vivo* tissue dynamics remain elusive. Here, using single-molecule analysis, endogenous gene-edited mRNAs and zebrafish *in-vivo* live-cell imaging, we report that mRNA polarisation acts as a molecular compass that orients motile cell polarity and spatially directs tissue movement. Clustering of protrusion-derived RNAseq datasets defined a core 192 bp localisation element underpinning precise mRNA targeting to incipient sites of filopodia formation at cell protrusions. Such targeting of the small GTPase, *RAB13*, generated tight spatial coupling of mRNA localisation, translation and protein activity, achieving precise subcellular compartmentalisation of RAB13 protein function to create a polarised domain of filopodia extension. Consequently, genomic excision of this localisation element and specific perturbation of endogenous *RAB13* targeting – but not translation – depolarised filopodial dynamics in motile endothelial cells and induced miss-patterning of nascent blood vessels *in-vivo*. Hence, mRNA polarisation, not expression, is the primary spatial determinant of the site of RAB13 action, preventing ectopic functionality at inappropriate subcellular loci and orienting tissue morphogenesis. Considering the unexpected spatial diversity of other polarised mRNA clusters we identified, mRNA-mediated compartmentalisation of protein function at distinct subcellular sites likely coordinates broad aspects of *in-vivo* tissue behaviour.

## RESULTS AND DISCUSSION

Dynamic subcellular polarisation of a myriad of proteins fundamentally shapes the front-rear polarity and directed movement of motile cells^4^. In parallel, cell migration is associated with subcellular polarisation of numerous mRNAs^1–3^. However, precise functional roles for targeted mRNAs remain unclear considering difficulties in identifying and manipulating endogenous transcript *cis* targeting elements^5^. Likewise, whether this phenomenon is relevant to modulation of tissue dynamics *in-vivo* remains an open question. To address these issues, we first aimed to define mRNA localisation motifs driving transcript polarisation in motile endothelial cells (ECs), as an initial step towards probing their function in coordinating blood vessel morphogenesis *in-vivo*. As a starting point, we identified 320 transcripts enriched in fractionated cellular protrusions of migrating primary human umbilical vein ECs (HUVECs) *in-vitro* (Fig.1a,b, Extended Data Table 1). *k*-means clustering analysis of these data alongside RNAseq datasets from unrelated cell types (NIH/3T3 fibroblasts^6^, MDA-MB231 metastatic breast cancer cells^7^, induced neuronal cells^8^) revealed unexpected cell type-specific diversity to transcript polarisation, with just five mRNAs exhibiting universal targeting to protrusions in all cell types tested (cluster *k*5; *RAB13*, *TRAK2*, *RASSF3*, *NET1*, *KIF1C*; Fig.1b, c). Strikingly, cluster *k*5 mRNAs all shared near-identical spatial distributions by single molecule FISH (smFISH)^9, 10^, being highly polarised to cellular protrusions relative to a control transcript, *GAPDH* (Fig. 1d-g). Moreover, *k*5 transcripts were highly spatially distinct from other clusters, such as mRNAs of cluster *k*7, which exclusively encoded secretory proteins^11–13^ and exhibited less-polarised peri-nuclear targeting (Fig.1g; Extended Data Fig.1b, e-f). Likewise, protrusion localisation of the well-established polarised mRNA, *ACTB*^14^, was significantly more diffuse than *k*5 mRNAs, as were other cluster *k*2 members (Fig.1g; Extended Data Fig.1b-d). Finally, protrusion-enriched mRNAs were also tightly clustered according to protein function (Extended Data Fig.1a), with *k*5 transcripts specifically encoding cell periphery-associated modulators of vesicle trafficking and membrane remodelling^15–19^. Hence, tight coupling of distinct spatial distributions of mRNAs with discrete protein functionalities indicated that the universal polarisation of cluster *k*5 mRNAs likely reflected a conserved functional requirement in motile cells.

**Figure 1.**
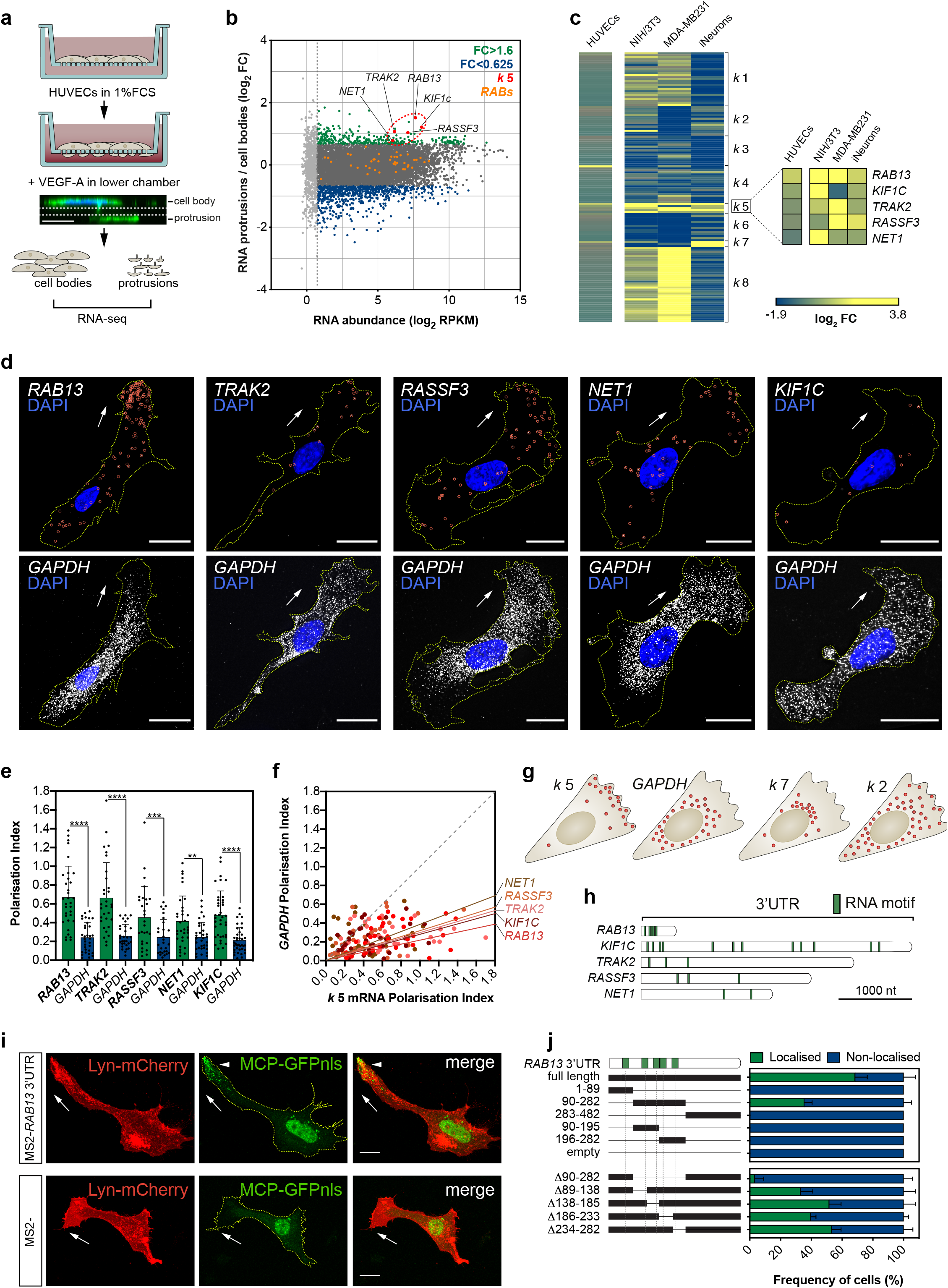
Clustering of RNAseq datasets defines a 192 bp localisation element driving mRNA targeting to protrusions. **a**, Strategy used in the RNAseq screen of mRNAs enriched in motile protrusions of HUVECs migrating through Transwell membranes. **b**, RNAseq data are plotted in reads per kilobase of transcripts per million reads (RPKM) against fold change (FC) levels of protrusions over cell bodies; light-grey data are transcripts with low expression and not considered; data are mean values represented in log_2_ (n=2 replicates). **c**, Heat map represents the *k* means clustering of transcript FC levels (protrusions over cell bodies) of other published extracted RNAseq datasets. The corresponding HUVEC FC levels are shown in parallel. **d**, smFISH co-detection of *k*5 mRNAs and *GAPDH* in exemplar HUVECs. **e**, Polarisation Index (PI) of *k*5 mRNAs and *GAPDH* in co-detected in HUVECs (n≥28 cells each co-hybridization). **f**, *k*5 mRNAs PIs plotted against respective *GAPDH* PIs. The slope of the coloured lines represents the average *k*5 mRNA/*GAPDH* PI ratio. **g**, Comparison of the mRNA distribution pattern of transcripts clustered in *k*2, *k5, k*7 and *GAPDH*. **h**, Diagram of *k*5 mRNA 3’UTRs and the positions of the RNA motif shared between transcripts. **i**, Representative bEnd.3 cells co-transfected with plasmids expressing Lyn-mCherry, MCP-GFPnls and 24xMS2-*RAB13* 3’UTR or 24xMS2. Arrowhead: non-nuclear localisation of MCP-GFPnls. **j**, Percentage of bEND.3 cells with MCP-GFP localised to protrusions when co-transfected with deletion versions of *RAB13* 3’UTR tagged with 24x MS2 hairpins; (n≥10 cells each triplicate). Data are mean±s.d. \*\**P*<0.01, \*\*\**P<*0.001, \*\*\*\**P*<0.0001. Arrows: orientation of RNA localisation. Scale bars: 20 μm.

Considering that the polarisation of cluster *k*5 mRNAs was particularly acute, highly stereotyped and uniquely conserved amongst cell types (Fig.1c-g), we hypothesised that these transcripts employed common targeting mechanisms. The *cis-* regulation of mRNA targeting is widely attributed to localisation elements (LE) within 3’ untranslated regions (3’UTR)^5, 20–24^. Indeed, despite identification of consensus *cis*- regulatory localisation elements being notoriously difficult, we detected repeat use of a conserved sequence motif in the 3’UTRs of all *k*5 transcript (Fig.1h; Extended Data Fig.2a). Moreover, this motif was striking in its clustering as five repeats within a short 3’UTR region of *RAB13*, a known polarised mRNA^23, 25^ (Fig. 1h). Potent localisation properties of this region were confirmed upon imaging of exogenous mRNAs in ECs using the MS2 system^26^ (Fig.1i,j; Extended Data Fig.2b). Using this approach, we identified a minimal 192 bp LE encompassing four motif repeats that was both necessary and sufficient to exclusively polarise mRNA at motile EC protrusions (Fig.1i,j). As *RAB13* was the only identified RAB small GTPase to exhibit such mRNA polarisation (Fig.1b), we hypothesised that this LE was critical for RAB13 function in motile cells. RAB13 is an established modulator of cortical F-actin crosslinking and cytoskeletal remodelling at leading front of migrating cells, via interaction with its effector protein, MICAL-L2^27–29^. Consistent with this function, live-cell imaging of *RAB13* 3’UTR dynamics revealed exclusive targeting of mRNA to sites of incipient filopodia formation in cellular protrusions, suggesting tight spatial coupling between *RAB13* mRNA localisation and RAB13 protein activity (Fig.2a-c; Supplementary Video1). Furthermore, quantification revealed a putative causal spatial relationship between *RAB13* proximity and filopodia frequency/stability (Fig.2b,c). Likewise, induction of cell migration drove a significant increase the levels and polarisation of *RAB13* mRNA *in-vitro* (Extended Data Fig.2c-e), further indicating dynamic involvement in leading-edge remodelling and establishment of cell polarity. Hence, these data revealed that polarisation of *RAB13* mRNA may spatially compartmentalise RAB13-mediated F-actin remodelling to orient motile cell polarity.

**Figure 2.**
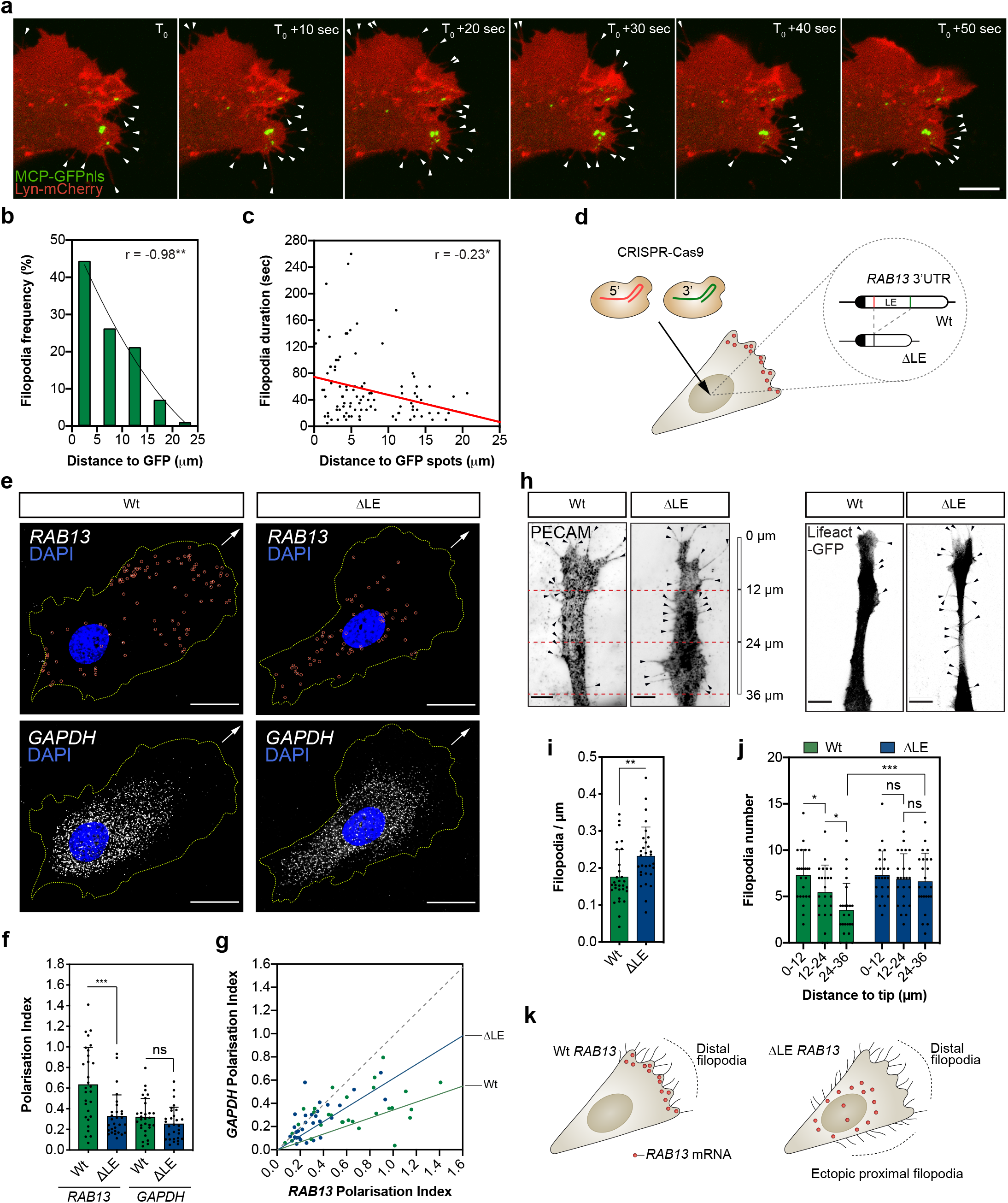
*RAB13* mRNA polarisation spatially orients filopodia dynamics. **a**, Representative time-lapse microscopy of a bEnd.3 cell co-transfected with plasmids expressing Lyn-mCherry, MCP-GFPnls and 24xMS2-*RAB13* 3’UTR. **b**, Frequency of newly formed filopodia formed within 5 μm intervals relative to the MCP-GFPnls signal (n=98 filopodia from 4 replicates). **c**, Distance to MCP-GFPnls signal of newly formed filopodia plotted against their duration. **d**, CRISPR-Cas9 strategy to excise the localisation element (LE) in the *RAB13* 3’UTR of HUVECs. **e**, smFISH co-detection of *RAB13* and control *GAPDH* in exemplar wild type (Wt) and ΔLE HUVECs. Arrows: orientation of RNA localisation. **f**, Polarisation Index (PI) of *RAB13* and *GAPDH* co-detected in Wt and ΔLE HUVECs (n=29 cells each genotype). **g**, *RAB13* PI plotted against respective *GAPDH* PI. The line slopes represent the average *RAB13*/*GAPDH* PI ratio. **h**, Exemplar Wt and ΔLE HUVECs co-cultured on fibroblast monolayers. **i**and **j**, Number of filopodia detected in co-cultured HUVECs per μm (**i**) and within 12 μm intervals relative to cell distal tip (**j**). **k**. Illustration of the spatial relationship between *RAB13* mRNA localisation and sites of filipodia production. Data are mean±s.d. \**P*<0.05, \*\**P*<0.01, *P*\*\*\*<0.001, ns: not significant (*P*>0.05). Arrowheads: filopodia. Scale bars: 6μm (**h**); 10 μm (**a**); 20 μm (**e**).

Studies probing the precise function of endogenous polarised mRNAs in motile cells are currently lacking, predominantly due to difficulties identifying *cis* targeting motifs and the potential propensity for genomic manipulation of mRNAs to perturb transcript stability and/or translation. However, precise genomic excision of the *RAB13* minimal LE in HUVECs using CRISPR-Cas9 tools did not perturb RAB13 protein expression (Extended Data Fig.3a-e), but did eradicate the polarised spatial pattern of *RAB13* localisation, such that transcript became diffusely distributed in ECs similar to *GAPDH* (Fig. 2e-g). Importantly, this loss of *RAB13* mRNA polarisation – but not loss of expression – was sufficient to depolarise filopodial dynamics in motile ECs (Fig.2h-k; Extended Data Fig.3f). When co-cultured with fibroblasts to mimic polarised blood vessel sprouting^30^, Wild type (Wt) HUVECs exhibited highly polarised filopodia extensions biased towards the leading edge of motile cell protrusions (Fig.2j). In contrast, filopodia in HUVECs lacking the *RAB13* LE were no longer spatially compartmentalised and were ectopically distributed homogeneously along the distal-proximal axis of migrating cells (Fig.2j). As a consequence, mutant ECs exhibited a significant increase in overall filopodia frequency (Fig.2i). Hence, this work revealed that tight control of *RAB13* mRNA localisation spatially specifies a polarised domain of filopodia extension in motile cells (Fig.2k).

These striking observations suggested that *RAB13* mRNA polarisation acts to exclusively spatially compartmentalise RAB13-mediated filopodia extension at distal sites. As such, targeting of *RAB13* mRNA and local translation could effectively block ectopic protein function at inappropriate subcellular loci to orient motile cells. However, this could only be achieved if the sites of *RAB13* mRNA localisation, translation and protein function were all tightly spatially coupled. Indeed, such coupling may be consistent with longstading proposals that newly translated RABs form a discrete protein pool from mature RABs, potentially with distinct interaction partners (e.g. specific RAB escorting proteins and GDP dissociation inhibitors) and separate biological functions^31–33^. Hence, local translation of polarised *RAB13* transcript may generate nascent protein with a distinct functionality to mature RAB13 at specific subcellular sites, thus achieving tight spatial compartmentalisation of RAB13-mediated membrane remodelling. As predicted, such coupling of polarised *RAB13* mRNA localisation with local translation was confirmed in EC protrusions upon detection of nascent protein using puromycinilation-proximity ligation assays (Puro-PLA)^34^.

HUVECs were cultured on Transwells and, prior to short incubation with puromycin, cell bodies removed to exclude detection of nascent puromycin-labelled proteins transported from the cell body to protrusions (Fig.3a). Isolated EC protrusions readily incorporated puromycin, which could be blocked upon pre-incubated with the translation inhibitor anisomycin (Extended Data Fig.4a), indicating active protein translation at the leading edge of migrating ECs. Importantly, Puro-PLA on isolated EC protrusions using antibodies recognising incorporated puromycin and endogenous RAB13 detected numerous distinct punctate of nascent synthesised RAB13, unlike anisomycin pre-treated and antibody-free controls (Fig.3a-c; Extended Data Fig.4b,c). Hence, polarised targeting of *RAB13* mRNA to motile cell protrusions drives local RAB13 translation. Moreover, spatial control of mRNA polarisation and local translation was coupled to regional compartmentalisation of RAB13 protein function, as loss of endogenous *RAB13* specifically disrupted filopodia dynamics only at sites of mRNA targeting (Fig.3e-g; Extended Data Fig.4d). Knockdown did not perturb *RAB13*- independent filopodia at proximal regions in ECs, but significantly depleted filopodia numbers at distal sites. This was not simply a consequence of spatial targeting of mature protein, as immunofluorescence assays revealed that RAB13 was homogeneously distributed throughout migrating cells (Extended Data Fig.4e). Alternatively, it was the location of *RAB13* mRNA itself that defined the domain of RAB13-dependent filopodia dynamics, as excision of the LE and diffuse miss-localisation of *RAB13* mRNA was sufficient to drive ectopic depolarised filopodia (Fig.2h-k). Hence, the site of *RAB13* mRNA localisation, translation and protein function are tightly spatially coupled in migrating cells. Thus, polarisation of *RAB13* transcript forms a molecular compass that achieves precise subcellular compartmentalisation of protein function, defines a polarised domain of filopodia extension and ultimately orients motile cell polarity (Fig.3g).

**Figure 3.**
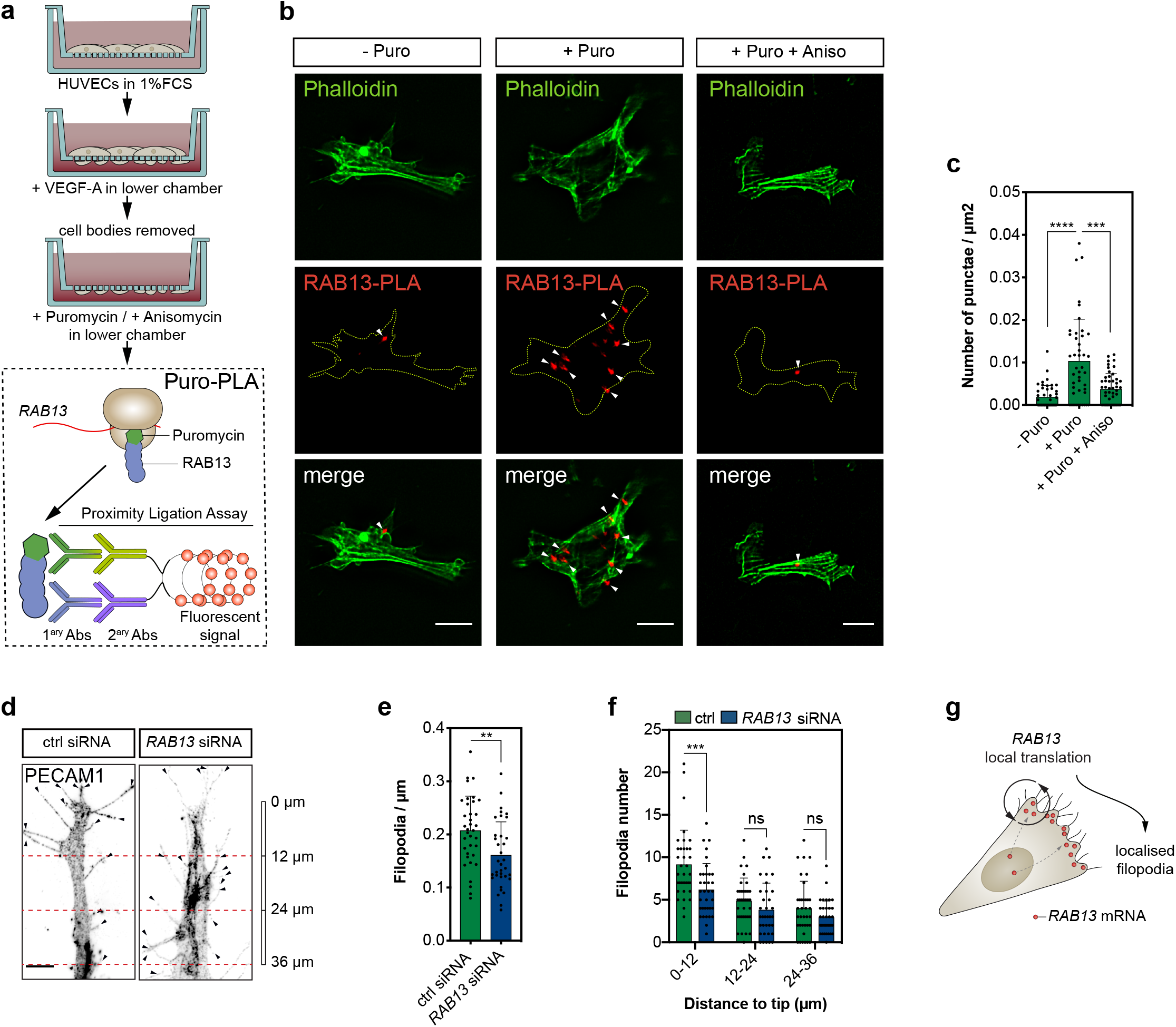
mRNA polarisation achieves spatial compartmentalisation of RAB13 translation and protein function. **a**, Strategy used to detect local protein synthesis in protrusions formed by HUVECs migrating through Transwell membranes. **b**, Representative Puro-PLA experiments detecting newly synthesised RAB13 in HUVEC protrusions present in the lower side of Transwell membranes. Arrowheads: Puro-PLA punctae. **c**, Quantification of RAB13 Puro-PLA punctae normalised to protrusion area; (n≥20 protrusions each duplicate). **d**, Exemplar control (ctrl) and *RAB13* siRNA-treated HUVECs co-cultured on fibroblast monolayers. **e**and **f**, Number of filopodia detected in co-cultured HUVECs per μm (**e**) and within 12 μm intervals relative to cell distal tip (**f)**. **g**, Illustration of the spatial relationship between the sites of *RAB13* mRNA localisation, local translation and RAB13 protein-mediated filopodia distribution. Data are mean±s.d. \*\**P*<0.01, \*\*\**P* <0.001, \*\*\*\**P*<0.0001, ns: not significant (*P*>0.05). Scale bars: 10 μm.

Although a widespread phenomenon, the functional role for localised mRNAs in tissue migration and vertebrate morphogenesis remains unexplored. Hence, having defined a key role for mRNA polarisation in the spatial control of EC behaviour *in-vitro*, we then sought to define the broader relevance of this phenomenon to modulation of tissue dynamics *in-vivo*. The production of polarised filopodial protrusions is a characteristic hallmark of motile endothelial tip cells, which lead new blood vessel branches during angiogenesis^35, 36^. As such, using live-cell imaging approaches in the zebrafish model system, we probed the function of *rab13* mRNA polarisation in the control of tip cell behaviour and angiogenesis *in-vivo*. Firstly, we generated a novel vascular-specific MCP-GFPnls transgenic strain, *Tg*(*fli1ep:MCP-GFPnls*), and monitored the dynamics of *rab13* 3’UTR mRNA targeting during intersegmental vessel (ISV) angiogenesis^36^ (Fig.4a; Extended Data Fig.5a and Supplementary Video2). Dynamic accumulation of MCP-GFPnls adjacent to and within filopodia at the leading edge of ISV tip cells revealed that *rab13* mRNA targeting *in-vivo* closely mirrored the 3’UTR-driven polarisation of *RAB13* mRNA *in-vitro*. Similarly, CRISPR-Cas9-mediated excision of a genomic fragment of the endogenous *rab13* 3’UTR locus (Δ3’UTR) confirmed that it also contained LEs critical for polarisation of zebrafish *rab13* mRNA (Fig.4b,c; Extended Data Fig.5b-e). In particular, smFISH for zebrafish *rab13* in explanted cells from dissociated *rab13*^*Δ3’UTR/Δ3’UTR*^ mutant embryos confirmed that mRNAs lacking these LEs were more diffusely distributed (Fig.4c), similar to observations in human ECs (Fig.2f). Hence, we reveal a previously unappreciated and conserved role for 3’UTR LEs in the dynamic polarisation of mRNA during cell migration *in-vivo*.

**Figure 4.**
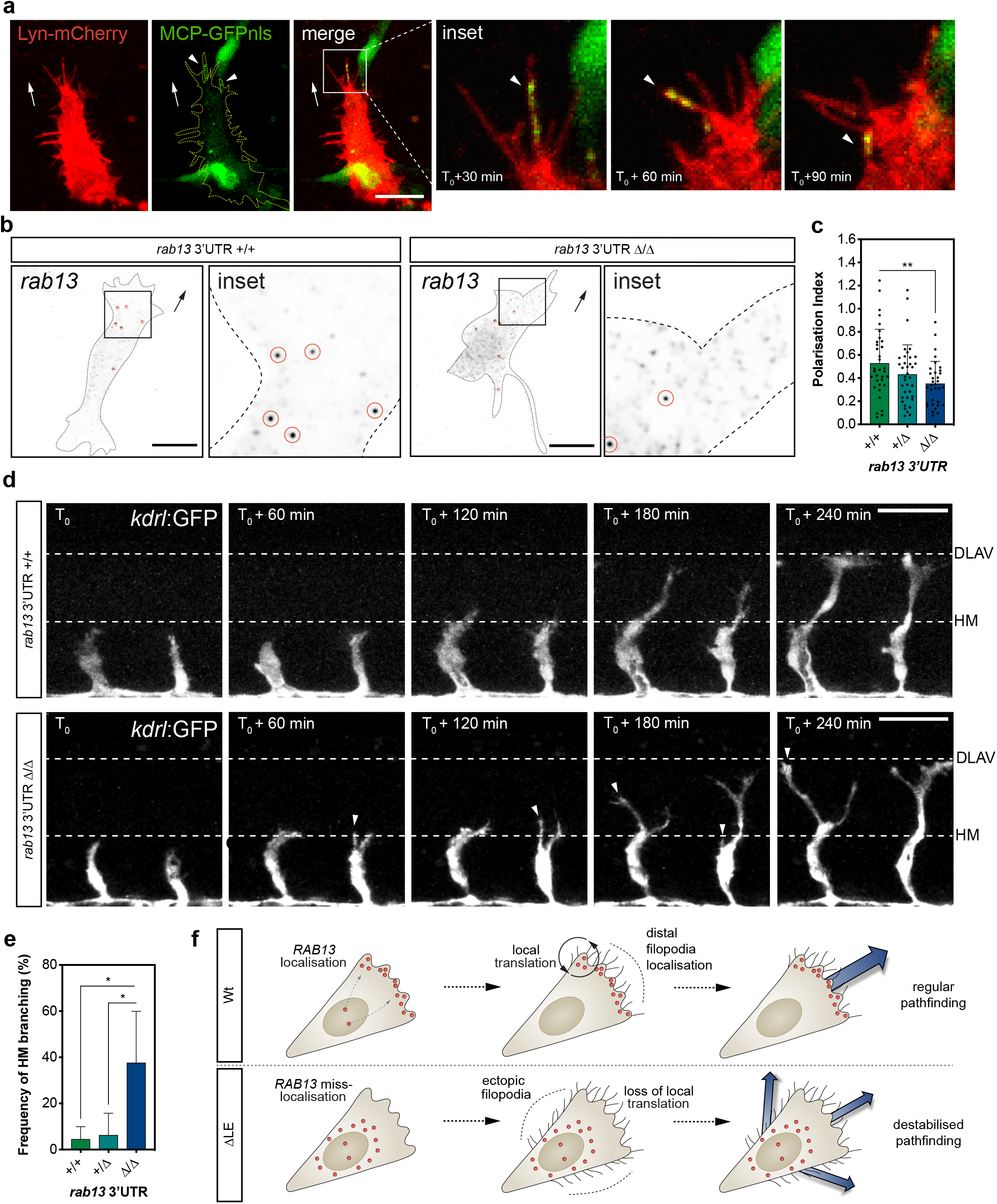
*rab13* mRNA polarisation orients blood vessel morphogenesis. **a**, Time-lapse microscopy of a representative *Tg(fli1ep:MCP-GFPnls)* (T_0_ = 28 hours post fertilisation (hpf)) ISV cell displaying mosaic expression of *rab13* 3’UTR tagged with Lyn-mCherry and 24xMS2 hairpins. Arrowheads: non-nuclear localisation of MCP-GFPnls. Arrows: direction of ISV sprouting. **b**, smFISH detection of *rab13* mRNA in 3’UTR +/+ and Δ/Δ cultured zebrafish embryo cells. **c**, Polarisation Index of *rab13* detected by smFISH in individual zebrafish cells (n≥33 cells each genotype). **d**, Time-lapse confocal microscopy of representative *rab13* 3’UTR +/+ and Δ/Δ *Tg(kdrl:EGFP*)^*s843*^ embryos starting at 25 hpf (T_0_). Arrowheads: extra branches emerging from the main ISVs at the HM position. **e**, Frequency of ISV ectopic branching occurring at the HM (n≥35 ISVs each genotype). **f**, Illustration of the role for *RAB13* mRNA localisation, local translation and compartmentalisation of RAB13 function in defining the orientation of EC filopodia dynamics, motile EC polarity and blood vessel pathfinding. *P*\*<0.05, \*\**P*<0.01. Data are mean±s.d. Scale bars: 10 μm (**b**), 50 μm (**a** and **d**).

During ISV branching, migrating tip cells must make key directional decisions, particularly when negotiating the multi-tissue junction of the horizontal myoseptum^37–39^ (HM; Extended Data Fig.5f). Hence, we hypothesised that mRNA localisation-mediated orientation of EC filopodia may indeed generate spatial cues that direct vascular tissue movement. Consistent with a key role for mRNA polarisation in the spatial coordination of vascular morphogenesis, live-cell imaging of ISVs branching in Wt and *rab13*^*Δ3’UTR/Δ3’UTR*^ embryos revealed that loss of *rab13* polarisation severely perturbed tip cell path-finding decisions (Fig.4d,e). Unlike ISVs in Wt and *rab13*^*+/Δ3’UTR*^ embryos that efficiently negotiated their way past the HM position, ISVs in *rab13*^*Δ3’UTR/Δ3’UTR*^ mutants struggled with this directional decision, resulting in a 7-fold increase in tip cells exhibiting ectopic miss-directed branches (Fig.4d,e). Importantly, *rab13* mRNA stability was unperturbed in *rab13*^*Δ3’UTR/Δ3’UTR*^ mutant embryos (Extended Data Fig.5g), consistent with excision of the *RAB13* LE *in-vitro* (Fig.2e), confirming that observed defects were not due to decreased *rab13* expression but were a consequence of perturbed mRNA localisation. Thus, we provide the first *in-vivo* evidence that spatial targeting of mRNAs and precise compartmentalisation of protein function generates key directional cues that orient motile cells and drive tissue morphogenesis (Fig.4f).

Here, using precise gene-editing of endogenous mRNAs, we reveal that tight spatial coupling of mRNA localisation, translation and protein function achieves the exclusive subcellular compartmentalisation of protein function. Hence, we define a novel paradigm for the spatial control of motile cell polarity and oriented tissue movement *in-vivo*. Taking into account that *RAB13* is one of only five mRNAs exhibiting conserved polarisation in all cell types tested, this function is likely a universally conserved mechanism for spatial coordination of complex morphogenetic events. Moreover, it is striking that all five cluster *k*5 mRNAs encode highly dynamic membrane trafficking and/or small GTPase-regulating proteins, all known to modulate cell motility. Hence, for classes of proteins normally constant in motion, mRNA-polarisation may be essential to spatially compartmentalise and precisely fix their site of function. Finally, this work reveals an unexpected spatial diversity to other identified clusters of polarised mRNAs. Considering our observations that the sites of mRNA targeting and protein function are tightly coupled, this raises the exciting possibility that other distinct mRNA distributions, such as the perinuclear localisation of cluster *k*7, reflect even broader functionalities for compartmentalised gene expression/function in the coordination of diverse aspects of tissue development and disease.

## METHODS

### Zebrafish husbandry

Zebrafish were grown and maintained according to UK Home Office regulation guidelines and all studies were approved by the University of Manchester Ethical Review Board.

### Embryo micro-injections and generation of zebrafish strains

To generate the transgenic zebrafish strain *Tg*(*fli1ep:MCP-GFPnls*) using Tol2 transposon transgenesis, 32 pg of Cerulean-H2B:bas*fli1ep*:MCP-GFPnls Tol2-based plasmid was co-injected with 32 pg Tol2 mRNA into one-cell stage AB zebrafish embryos. The next day, embryos with mosaic GFP expression were selected, raised into adulthood and then outbred to AB zebrafish to identify founders with germ-line transmission of the transgene. Adult *Tg*(*fli1ep:MCP-GFPnls*) were inbred and one-cell stage embryos were co-injected with 32 pg of Cerulean-H2B:bas*fli1ep:*Lyn-mCherry-24xMS2-*rab13*-3’UTR Tol2-based plasmid and 32 pg Tol2 mRNA for mosaic expression analysis.

The mutant *rab13* Δ3’UTR strain was generated with CRISPR-Cas9 tools. One-cell stage *Tg*(*kdrl:EGFP*)^*s843*^ embryos^40^ were injected with 150 pg of each *in vitro* transcribed gRNA and co-injected with 150 pg Cas9 NLS nuclease (New England Biolabs). Embryos were raised to adulthood and outbred to AB zebrafish to identify founders with germ-line transmission deletions in the *rab13* 3’UTR. Heterozygous animals harbouring a 482-nucleotide deletion in the *rab13* 3’UTR (*Tg*(*kdrl:EGFP*)^*s843*^ *rab13*^+/Δ3’UTR^) were in-crossed and the resulting embryos were used for live-cell imaging analysis.

### gRNA generation and *in vitro* transcription

The online CRISPRscan tool^41^ was used to design gRNAs targeting the zebrafish 3’UTR region in the *rab13* locus (Extended Data Table 2). Next, 0.3 μM oligonucleotides comprising the target sequences (flanked by the T7 promoter and the Tail annealing sequence) were mixed with 0.45 μM Tail primer (Extended Data Table 2) and PCR amplified with Platinum Pfx DNA polymerase (ThermoFisher Scientific) in a T100 thermal cycler (BioRad). The following cycling conditions were used: 1 cycle of initial denaturation at 94 °C for 10 minutes, 30 cycles of denaturation at 94 °C for 30 seconds, annealing at 45 °C for 30 seconds, extension at 68 °C for 30 seconds, and a final extension cycle at 68 °C for 7 minutes. Subsequently, 200 ng of PCR amplified templates were used to transcribe gRNAs using a MEGAshortscript T7 Transcription Kit (ThermoFisher Scientific), following the manufacturer’s recommendations. To synthesise Tol2 mRNA, 1 μg NotI-linearised pCS2-TP plasmid was transcribed using a SP6 mMESSAGE mMACHINE kit (ThermoFisher Scientific) according to manufacturer’s protocol.

### Embryo genotyping

Genomic DNA was extracted by incubating either whole embryos or embryo heads in lysis buffer (10 mM Tris HCl pH8, 1 mM EDTA, 80 mM KCl, 0.3 % NP40, 0.3 % Tween) containing 0.5 μg/μl Proteinase K (Promega) at 55 °C for 1-2 hours, followed by a denaturation step at 95 °C for 15 minutes in a T100 thermal cycler. Genotyping PCR was performed using 2 μl genomic DNA, 0.4 μM zebrafish genotyping primers (Extended Data Fig.5A, Table 2) and 1 x MyTaq Red DNA Polymerase (Bioline) according to the manufacturer’s protocol in a T100 thermal cycler. PCR reactions were resolved in 1 % agarose (Bioline) gels containing 0.5 μg/ml Ethidium Bromide (Sigma) for analysis. PCR products were cloned into TOPO-TA vectors (ThermoFisher Scientific) according to the manufacturer’s protocol and analysed via Sanger sequencing on an ABI 3730 device.

### Cell culture, scratch wound and co-culture angiogenesis assays

Trunks of 26-28 hpf embryos were incubated in Trypsin-EDTA solution (Sigma) at 28 °C for 15 minutes. Trypsinisation was quenched with complete L-15 medium (Sigma) containing 10 % Fetal Bovine Serum (FBS, Sigma) and 10 U/ml-100μg/ml Penicillin-Streptomycin (Sigma). Cells were pelleted at 2000 rpm for 5 minutes at room temperature (RT), resuspended in complete L-15 medium, plated on Laminin-coated (Sigma) coverslips in 24 well plates and maintained at 28 °C for 18 hours. HUVECs (PromoCell) were cultured in complete ECGM2 (PromoCell) in gelatin-coated (Millipore) dishes. Human Pulmonary Fibroblasts (HPF; PromoCell) were cultured in M199 (ThermoFisher Scientific) containing 10 % FBS, 50 μg/ml Gentamycin (Sigma) and 50 ng/ml Amphotericin (Sigma). Brain endothelial (bEnd.3) were cultured in DMEM (Sigma) supplemented with 10 % FBS, 10 ng/ml Recombinant Human VEGF-A (PeproTech) and 10 U/ml-100μg/ml Penicillin-Streptomycin. For scratch wound assays, HUVECs cultured on gelatin coated coverslips were grown to confluence and used in scratch wound assays as described elsewhere^42^. Co-cultures of HUVEC and HPF and the corresponding siRNA-mediated knockdown experiments were performed as previously described by Hetheridge *et al*.^30^.

### CRISPR-Cas9 cell editing and cell transfections

To edit RAB13 locus, HUVECs were transfected with Alt-R CRISPR-Cas9 ribonucleoprotein complexes (Integrated DNA Technologies) targeting the 90-282nt localisation element within the 3’UTR. Briefly, each sequence-specific crRNA (Extended Data Table 2) was mixed with tracrRNA at 1:1 50 μM, incubated at 95 °C for 5 minutes in a T100 thermal cycler and allowed to cool to RT for 60 minutes. Next, 12 μM each crRNA:tracrRNA (gRNA) was incubated with 20 μM Alt-R Cas9 nuclease in PBS (Sigma) at RT for 20 minutes to form ribonucleoprotein complexes and mixed with 500×10^3^ HUVECs. Additionally, 2 μg pmaxGFP Vector (Lonza) was included in the HUVEC-ribonucleoprotein mix to identify transfected cells. Transfections were performed in a Nucleofector 2b Device (Lonza), using a HUVEC Nucleofector kit (Lonza) according to manufacturer’s instructions and the cells were further cultured for 72 hours. Afterwards, single GFP-expressing cells were isolated in a FACS Aria Fusion cell sorter (BD Biosciences) into gelatin-coated 96 well plates to grow individual clones. Genomic DNA was extracted from expanded HUVEC clones and PCR-analysed with sequence-specific primers (Extended Data Fig.3A, Table 2) as described for zebrafish embryo genotyping.

Knockdown experiments were performed with ON-TARGETplus Non-targeting control or *RAB13* siRNAs (Horizon) using GeneFECTOR (VennNova) as previously described^30^.

For *in vitro* MS2 experiments, bEnd.3 cells were transfected with pcDNA3-Lyn-mCherry, pCS2-MCP-GFPnls and different versions of pcDNA3-*HBB*-24XMS2SL-*RAB13* 3’UTR. Briefly, 100×10^3^ cells / well cultured in 6 well plates were transfected with 0.8 μg each plasmid DNA using Lipofectamin2000 following the manufacturer’s protocol (ThermoFisher Scientific) and analysed 48 hours later.

### Transwell assays and cell body / protrusion fractionation

Transwell experiments to segregate cell bodies and protrusions were performed as described elsewhere^23^, with the following modifications: 1.5×10^6^ HUVECs were cultured for 2 hours in 24 mm Transwells (Costar), containing 3μm-pore polycarbonate membranes, in M199 (ThermoFisher Scientific) supplemented with 1 % FBS. Subsequently, 25ng/ml VEGF-A was added to the lower chambers to promote cell migration over the next hour. While only 1 Transwell was used for the cell body fraction, 2 Transwells were used to harvest each HUVEC protrusion sample.

### RNA isolation, qPCR and RNAseq

Embryo and cell-derived RNA was isolated using a RNAqueous-Micro kit (ThermoFisher Scientific) according to the manufacturer’s protocol. For gene expression analysis, cDNA was synthesised with a High-Capacity RNA-to-cDNA kit (ThermoFisher Scientific) following the manufacturer’s protocol. qPCR experiments were performed with 1-2 μl cDNA, 0.25 μM gene-specific primers (Extended Data Table 2) and 1 x Power SYBR Green Master Mix (ThermoFisher Scientific) in a StepOne Real-Time PCR System (Applied Biosystems). *GAPDH* expression was used to normalise gene expression levels and the relative mRNA levels were analysed with the *2*^-Δ;Δ;*CT*^ method. For RNAseq, quality and integrity of RNA samples obtained from HUVEC cell bodies and protrusions were assessed using a 2200 TapeStation (Agilent Technologies). Next, RNAseq libraries were generated using the TruSeq Stranded mRNA assay (Illumina) according to the manufacturer’s protocol. Adapter indices were used to multiplex libraries, which were pooled prior to cluster generation using a cBot instrument. The loaded flow-cell was then paired-end sequenced (76 + 76 cycles, plus indices) on an Illumina HiSeq4000 instrument. Finally, the output data was demultiplexed (allowing one mismatch) and BCL-to-Fastq conversion performed using Illumina’s bcl2fastq software, version 2.17.1.14.

### smFISH

Zebrafish cells and HUVECs cultured on either Laminin or gelatin-coated coverslips, respectively, were fixed in methanol free 4 % formaldehyde (ThermoFisher Scientific) and used in smFISH assays. Briefly, cells were permeabilised with 70 % Ethanol at RT for 1 hour or 4 °C overnight, washed with smFISH wash buffer (2 X SSC, 10 % formamide) and incubated with smFISH probes (Extended Data Table 3) in smFISH hybridisation buffer (10 % dextran sulfate, 2 X SSC, 10 % formamide) at 37 °C overnight. Afterwards, cells were washed with smFISH wash buffer twice at 37 °C for 30 minutes, washed once with 2 X SSC for 10 minutes, counterstaining with 1 μg/ml DAPI (Sigma) and washed twice with PBS for 5 minutes at RT. Coverslips were air-dried and mounted on microscope slides with ProLong Gold Antifade Mountant (ThermoFisher Scientific). All probes targeting protrusion-enriched mRNAs were designed with Stellaris Probe Designer (LGC Biosearch Technologies), synthesised and labelled with Quasar 570 (LGC Biosearch Technologies) or Alexa 594 fluorophores (Integrated DNA Technologies). Co-hybridisation experiments were carried out with predesigned *GAPDH* probes labelled with Quasar 670 (LGC Biosearch Technologies).

### Puro-PLA and immunofluorescence (IF)

For Puro-PLA, cell bodies of HUVECs cultured in Transwells were scraped off and remaining protrusions were exposed to 3 μM Puromycin (Sigma) added to lower chambers for 6 minutes. In translation inhibition experiments, 40 μM Anisomycin (Sigma) was added to the lower Transwell chamber 30 minutes before cell body removal and 6 minutes after cell body removal together with 3 μM Puromycin. Subsequently, HUVEC protrusions grown in Transwell membranes were fixed in methanol free 4 % formaldehyde, removed from the Transwell inserts and used in Puro-PLA experiments as described elsewhere^34^. Following the Puro-PLA protocol, Transwell membranes were incubated for 20 minutes with 1:40 Alexa Fluor 488 Phalloidin (ThermoFisher Scientific) in PBS, washed in Duolink wash buffer B (Sigma) and mounted on microscope slides with Duolink *In Situ* Mounting Medium containing DAPI (Sigma).

For IF experiments, cells and Transwell membranes containing protrusions were permeabilised in PBS containing 0.2-0.5 % Triton-X100 (Sigma), blocked in 4 % goat serum (Sigma) for 15 minutes, and incubated with primary antibodies in blocking solution at 4 °C overnight. Next, cells were washed in PBS containing 0.2 % Tween, incubated with secondary antibodies at RT for 1 hour, counterstaining with 1 μg/ml DAPI and washed again. Transwell membranes were further incubated with 1:40 Phalloidin Alexa Fluor 488 (ThermoFisher Scientific) in PBS at RT for 20 minutes before washing. Cells and Transwell membranes were mounted with ProLong Gold Antifade Mountant (ThermoFisher Scientific).

### Western blotting

Proteins were extracted with RIPA buffer (25 mM Tris-HCl pH 7.6, 150 mM NaCl, 1 % NP-40, 1 % sodium deoxycholate, and 0.1 % SDS) and quantified with Pierce BCA protein assay kit (ThermoFisher Scientific) following the supplier’s recommendations. Samples were denatured with Laemmli buffer (250 mM Tris-HCl pH 6.8, 2 % SDS, 10 % glycerol, 0.0025 % bromophenol blue, 2.5 % β-mercaptoethanol) at 95 °C for 5 minutes, loaded on 10% Mini-PROTEAN TGX precast protein gels (Bio-Rad) and separated in a Mini-PROTEAN Electrophoresis System (Bio-Rad). Proteins were transferred onto nitrocellulose membranes using a Trans-Blot Turbo Transfer System RTA kit following the manufacturer’s protocols (Bio-Rad). Subsequently, membranes were blocked in 5 % milk (Sigma) or 5 % BSA (Sigma) in TBS containing 0.1% Tween at RT for 1 hour and incubated with primary antibodies at 4 °C overnight. The next day membranes were washed with TBS containing 0.1% Tween, incubated with secondary antibodies at RT for 1 hour and washed again. Signal detection was carried out with SuperSignal West Dura Extended Duration Substrate (ThermoFisher Scientific) according to the supplier’s recommendations.

### Antibodies

Primary and secondary antibodies were used at the following concentrations: 1:1600 mouse PECAM-1 89C2 (Cell Signaling Technology), 1:100 rabbit RAB13 (Puro-PLA and IF, Millipore), 1:1000 rabbit RAB13 (Western blotting, Cambridge Bioscience), 1:3500 mouse Puromycin (Kerafast), 1:1000 rabbit β-Tubulin 9F3 (Cell Signaling Technology), 1:200 mouse ZO-1 1A12 (ThermoFisher Scientific), 1:500 goat anti-mouse Alexa Fluor 488 or Alexa Fluor 568, (ThermoFisher Scientific) and 1:5000 goat anti-mouse HRP-linked antibody (Cell Signaling Technology).

### Plasmid construction

The pCS2-MCP-GFPnls plasmid used in *in vitro* MS2-system assays was generated excising a MCP-GFPnls fragment with SpeI and KpnI from pMS2-GFP, a gift from Robert Singer (Addgene plasmid # 27121)^43^, and subcloning it into a pCS2+ vector using the XbaI and KpnI sites.

To construct the Cerulean-H2B:*basfli1ep:*MCP-GFPnls Tol2-based plasmid for *in vivo* studies, MCP-GFPnls was amplified from pMS2-GFP with sequence 0.3 μM specific primers (Extended Data Table 2) and Platinum Pfx DNA polymerase in a T100 thermal cycler. Subsequently, the PCR product was cloned into a pDONR221 P3-P2 using Gateway Technology (ThermoFisher Scientific) according to manufacturer’s manual. The final Tol2-based construct was assembled into the pTol2Dest(R1R2) (Addgene plasmid # 73484)^44^ using Gateway 3-fragment recombination with pE(L1L4)Cerulean-H2B in the first position, pE(R4R3)*basfli1ep*^7^ in the second position and pE(L3L2)MCP-GFPnls in the third position.

For *in vitro* MS2-system experiments, the 3’UTR of human *RAB13* was PCR amplified with 0.3 μM sequence-specific primers (Extended Data Table 2) using Platinum Pfx DNA polymerase in a T100 thermal cycler and the resulting PCR product was cloned using a Zero Blunt PCR cloning kit (ThermoFisher Scientific), following the manufacturer’s manual. Next, the human *HBB* gene was PCR amplified using 0.3 μM sequence-specific primers (Extended Data Table 2) and Platinum Pfx DNA polymerase in a T100 thermal cycler and cloned into the NotI and BamHI sites of the pCR4-24XMS2SL-stable plasmid, a gift from Robert Singer (Addgene plasmid # 31865)^26^. Subsequently, a multiple cloning site (MCS, Extended Data Table 2) was introduced into the BglII and SpeI sites of pCR4-*HBB*-24XMS2SL and the recombinant *HBB*- 24XMS2SL-MCS sequence was subcloned into the pcDNA3 mammalian expression vector (ThermoFisher Scientific) using the NotI and XbaI sites. The full-length 482nt *RAB13* 3’UTR was then sub-cloned into pcDNA3-*HBB*-24XMS2SL-MCS using NheI and XhoI sites. Alternatively, truncated and deletion versions of *RAB13* 3’UTR were generated by PCR using 0.3 μM sequence-specific primers (Extended Data Table 2) and Platinum Pfx DNA polymerase or using QuikChange II Site-Directed Mutagenesis kit (Agilent Technologies) following the manufacturer’s instructions and introduced into the pcDNA3-*HBB*-24XMS2SL-MCS using the NheI and XhoI sites.

In order to generate the zebrafish MS2-system reporter construct, the 24XMS2SL cassette was firstly subcloned from pCR4-24XMS2SL-stable into a *kdrl:*Lyn-mCherry Tol2 based plasmid^45^ using a BamHI site. Next, the zebrafish *rab13* 3’UTR was PCR amplified with 0.4 μM sequence specific primers (Extended Data Table 2) and MyTaq Red DNA Polymerase from zebrafish genomic DNA in a T100 thermal cycler and then subcloned into the Tol2 *kdrl:*Lyn-mCherry-24XMS2SL plasmid using NheI and BglII sites. The resulting Lyn-mCherry-24XMS2SL-*rab13* 3’UTR recombinant sequence was amplified with 0.3 μM sequence-specific primers (Extended Data Table 2) and Platinum Pfx DNA polymerase in a T100 thermal cycler and subcloned into a pDONR221 P3-P2 using Gateway Technology. Lastly, the final Tol2-based construct was assembled into the pTol2Dest(R1R2) using Gateway 3-fragment recombination with pE(L1L4)Cerulean-H2B in the first position, pE(R4R3)*basfli1ep* in the second position and Lyn-mCherry-24XMS2SL-*rab13* 3’UTR in the third position.

All plasmid maps and details are available upon request.

### Microscopy

Confocal time-lapse imaging of zebrafish embryos was carried out as previously described^45^. MS2 system-transfected cells were live imaged every 5 seconds in a Nikon A1R inverted confocal microscope equipped with an Okolab incubation chamber, using a 60 X objective. Fixed images of cultured cells and Transwell membranes were acquired on an Olympus IX83 inverted microscope using Lumencor LED excitation, either a 60 X/ 1.42 PlanApo or a 100 X/ 1.35 UplanApo objective and a Sedat QUAD (DAPI/FITC/TRITC/Cy5) filter set (Chroma 89000). The images were collected using a R6 (Qimaging) CCD camera with a Z optical spacing of 0.2 μm. Raw images were then deconvolved using the Huygens Pro software (SVI) and maximum intensity projections of these images were used for analysis.

### smFISH spot quantification, Polarisation Index (PI) and filopodia analysis

Processed smFISH images were used to calculate mRNA polarisation with the PI metric developed by Park *et al*.46 and to assess mRNA spot number with FISHQuant47.

For the studies of filopodia distance to GFP signal in MS2-system experiments, filopodia parameters (position, duration, and frequency) of MS2-system transfected cells were determined using Filopodyan plugin for FIJI^48^. Subsequently, the coordinates of GFP particles were extracted with the TrackMate plugin for FIJI^49^ and the Euclidean distance between the base of newly formed filopodia and the nearest GFP particle was calculated.

### Statistics

All data are represented as mean±standard deviation. Statistical analysis of the data was carried out using GraphPad prism software. Differences in smFISH spot, Puro-PLA and filopodia numbers, protein and RNA levels, ISV length and branching were interrogated using t-tests when the data were normally distributed or Mann-Whitney tests when the data did not pass normality tests. The correlation between GFP distance and filopodia frequency or duration were assessed with Spearman correlation tests. Statistical significance is reported for *P*<0.05.

*k*-means clustering of RNAseq data was performed using Morpheus (Broad Institute, https://software.broadinstitute.org/morpheus/index.html). The number of clusters was defined by the number of cell types and by the possible transcript statuses (enriched or depleted) - 2^3^ =8. Gene Ontology studies were performed using DAVID^50, 51^.

## ACKNOWLEDGEMENTS

We wish to thank members of the University of Manchester Biological Services, Genomic Technologies, Bioimaging Facilities and Flow Cytometry Facilities, for technical support. We are grateful to E. Schuman and S. tom Dieck at the MPI for Brain Research, Frankfurt, Germany, for help and guidance setting up Puro-PLA assays. We also thank N. Papalopulu and group members at the University of Manchester, UK, for critical feedback, reagents and materials. This work was funded by the Wellcome Trust (095718/Z/11/Z to S.P.H.), Wellcome Institutional Strategic Support Fund (7064646 to G.C.) and the British Heart Foundation (PG/16/2/31863 to S.P.H.).

## AUTHOR CONTRIBUTIONS

Conceptualisation, G.C. and S.P.H.; Methodology, G.C.; Formal Analysis, G.C.; Investigation, G.C., J.B. and N.T.; Writing – Original Draft, G.C. and S.P.H.; Supervision, G.C. and S.P.H.; Funding acquisition, G.C. and S.P.H.

## COMPETING INTERESTS

The authors declare no competing interests

## ADDITIONAL INFORMATION

Correspondence and requests for materials should be addressed to S.P.H. or G.C.

