## Extended Data for "mRNA compartmentalisation spatially orients tissue morphogenesis"

**Extended Data Figure 1. Clustering of RNAseq datasets defines unexpected cell type specific diversity to mRNA polarisation. a,** Gene ontology analysis of the mRNAs enriched across cell-protrusion types and grouped by *k* means clustering (Fig. 1c). **b,** Detail of the heat map shown in Fig. 1c representing fold change (FC) levels (protrusions over cell bodies) of mRNAs present in clusters *k* 2 an*d k* 7. The corresponding HUVEC FC levels are shown in parallel. **c** and **e,** smFISH co-detection of *k* 2 (**c**) and *k* 7 (**e**) mRNAs and *GAPDH* in exemplar HUVECs. Arrows: orientation of RNA localisation. **d** and **f,** Polarisation Index of *k* 2 (**d**) and *k* 5 (**f**) mRNAs and *GAPDH* co-detected in HUVECs (n≥25 cells each co-hybridisation). Data are mean±s.d. *$P$<0.05, **$P$<0.01, ***$P$<0.001, ns: not significant ($P$>0.05). Scale bars: 20 μm.

**Extended Data Figure 2. Visualisation of the dynamics of *RAB13* mRNA polarisation**. **a,** RNA motif over-represented in *k* 5 mRNA 3'UTRs (Fig. 1h). **b,** CMV promoter-driven expression of MCP-GFPnls and *hHBB*-24xMS2-tagged *RAB13* 3'UTR. The visualisation of MCP-GFPnls bound to 24xMS2 allows the identification of the minimal region in the 3'UTR of *RAB13* necessary for its localisation (Fig. 1i,j). **c,** Scratch wound assay generates a free edge on a confluent monolayer of HUVECs and encourages cell migration (top). smFISH co-detection of *RAB13* mRNA and *GAPDH* mRNA in representative HUVECs migrating in a scratch wound assay. ZO-1 immunolabeling defines cell boundaries (bottom). Arrows: direction of migration. **d,** Polarisation Index (PI) of *RAB13* and *GAPDH* co-detected in exemplar HUVECs cultured in a scratch wound assay (n≥28 cells each group). **e,** Quantification of the number of *RAB13* mRNA smFISH spots per cell. Data are mean±s.d. *$P<0.05$, ***$P<0.001$, ns: not significant ($P>0.05$). Scale bars: 20 μm.

**Extended Data Figure 3. CRISPR-Cas9 editing of the *RAB13* 3'UTR does not alter protein levels. a,** CRISPR-Cas9 strategy to excise the localisation element (LE) in the *RAB13* 3'UTR of HUVECs. The wild type (Wt) *RAB13* exon 8 is represented with its coding sequence in dark and the 3'UTR in clear boxes. The 5' and 3' gRNA-targeted regions are represented with red and green lines. Arrows: relative positions of the forward (F) and reverse (R) PCR primers used to identify HUVECs with CRISPR-Cas9-mediated excision (Δ) of the LE. **b,** Representative genotyping PCR demonstrates the band size shift in ΔLE HUVECs. **c,** Chromatogram confirming the excision of the LE within *RAB13* 3'UTR. **d,** Detailed DNA sequence depicting nucleotide positions within the *RAB13* 3'UTR of Wt and the exemplar ΔLE HUVECs (**c**). PAM sequences are represented with dark red boxes. gRNA-targeted sequences are represented with light red boxes. **e,** Representative Western blotting (WB) of Wt and ΔLE HUVECs (left). Densitometry analysis of WB data (triplicates each genotype) (right). **f,** Number of filopodia detected in individual clones of co-cultured HUVECs within 12 μm intervals relative to cell distal tip. Data are mean±s.d. ns: not significant.

**Extended Data Figure 4. Protein translation in endothelial cell protrusions. a,** Immunofluorescence (IF) analysis of HUVEC protrusions generated on the underside of Transwell membranes and exposed to Puromycin (n=2 replicates). **b,** Representative Puro-PLA experiments detecting of newly synthesised RAB13 in HUVEC protrusions present in the lower Transwell membrane side. Arrowheads: Puro-PLA punctae. **c,** Quantification of RAB13 Puro-PLA punctae normalised to protrusion area; (n≥20 protrusions each duplicate). **d,** Representative Western blotting (WB) of siRNA transfected HUVECs (left). Densitometry analysis of WB data (triplicate experiments) (right). e. Representatiive RAB13 IF assay (left) and without (right) primary antibody on migrating HUVECs. Data are mean±s.d. ***P*<0.01, *****P*<0.0001. Scale bars: 10 µm.

**Extended Data Figure 5. Loss of *rab13* mRNA polarisation in CRISPR-Cas9-edited zebrafish cells. a,** *Tg(fli1ep:MCP-GFPnls)* zebrafish embryo 26 hpf displays vascular-specific expression of MCP-GFPnls. Inset shows the nuclear expression of MCP-GFPnls in dorsal aorta (DA) and in sprouting intersomitic vessels (ISVs) reaching the horizontal myoseptum (HM) (left). *fli1* enhancer/promoter (*fli1ep*)-driven expression of reporter constructs. Simultaneous translation of Lyn-mCherry reporter and binding of MCP-GFPnls to 24xMS2-*rab13* 3'UTR. (right). **b,** CRISPR-Cas9 strategy to excise a nucleotide fragment from the zebrafish *rab13* 3'UTR. The wild type (Wt) *rab13* exon 8 is represented with its coding sequence in dark and the 3'UTR in clear boxes; the 5' and 3' gRNA-targeted regions are represented with red and green lines, respectively. Arrows: relative positions of the forward (F) and reverse (R) PCR primers used to identify animals with CRISPR-Cas9-mediated deletions (Δ) in the *rab13* 3'UTR. **c,** Representative genotyping PCR demonstrates the band size shift in zebrafish harboring a Δ482 *rab13* 3'UTR. Asterisk marks a heteroduplex formed between Wt and Δ482 *rab13* 3'UTR PCR amplicons. **d,** Chromatogram confirming the CRISPR-Cas9-mediated generation of Δ482 *rab13* 3'UTR. **e,** Detailed DNA sequence depicting nucleotide positions within the Wt and Δ482 *rab13* 3'UTR. **f,** Scheme depicts stages of zebrafish ISV sprouting. DA: dorsal aorta. DLAV: dorsal longitudinal anastomotic vessel. HM: horizontal myoseptum. NC: notochord. NT: neural tube. **g.** *rab13* mRNA levels in individual 26-28 hpf clutch-matched sibling embryos (n≥9 embryos each genotype). Data are mean±s.d. ns: not significant. Scale bar: 20 μm.

**Extended Data Table 2. Oligonucleotide sequences related to Methods.**

**CRISPr/Cas9**

| **Name** | **Sequences (5' – 3')** |
|---|---|
| gRNA 5' *rab13* 3' UTR | **taatacgactcactata**GGAGCCTGTCCTCTGGAAAAgttttagagctagaa |
| gRNA 3' *rab13* 3' UTR | **taatacgactcactata**GGGGATGGATGCAAGAGTTAgttttagagctagaa |
| gRNA Tail | AAAAGCACCGACTCGGTGCCACTTTTTCAAGTTGATAACGGACT AGCCTTATTTTAACTTGCTATttctagctctaaaac |
| 5' crRNA *RAB13* 3' UTR | AAATAGCAGAGGGGCTTGGA |
| 3' crRNA *RAB13* 3' UTR | TCAGGCTTCAGACCTTACCT |

**Genotyping**

| **Name** | **Sequences (5' – 3')** |
|---|---|
| *rab13* 3' UTR F | AATTTTGCTGTGACGAGCCA |
| *rab13* 3' UTR R | TGGACCAAATGCACACACAA |
| *RAB13* 3' UTR F | CTGAGGACCCTTTCTTGCCT |
| *RAB13* 3' UTR R | TTGTGCAAATGGTGGCCTTT |

**qPCR**

| **Name** | **Sequences (5' – 3')** |
|---|---|
| *rab13* F | GCATACTACAGAGGGGCCAT |
| *rab13* R | CATTCGACTTACACCCGCTG |
| *gapdh* F | GTGGAGTCTACTGGTGTCTTC |
| *gapdh* R | GTGCAGGAGGCATTGCTTACA |

**Cloning**

| **Name** | **Sequences (5' – 3')** |
|---|---|
| NheI *rab13* 3' UTR F | GGAAGCTAGCGCGAACCATTTTCCAGAGG |
| BamHI *rab13* 3' UTR R | GGAAGGATCCTTTAACATTTTCTAATTTATTC |
| *att*B3 MCP-GFPnls F | ggggacaactttgtataataaagttgGCCACCATGGGCTACCCCTACG |
| *att*B2 MCP-GFPnls R | ggggaccactttgtacaagaaagctgggtTTATACCTTTCTCTTCTTTTTGG |
| *att*B3 Lyn-mCherry F | ggggacaactttgtataataaagttgGCCACCATGGGCTGCATCAAG |
| *att*B2 *rab13* 3' UTR R | ggggaccactttgtacaagaaagctgggtTGGCCGTATCTTCGCAGATC |
| multiple cloning site F | GATCTATCGATGGTACCGCTAGCGATATCCTCGAGA |
| multiple cloning site R | TAGTCTCGAGGATATCGCTAGCGGTACCATCGATA |
| NotI *HBB* F | GGAAGCGGCCGCACATTTGCTTCTGACACAACTG |
| BglII *HBB* R | GGAAAGATCTCCCAGTTTAGTAGTTGGAC |
| XbaI 1nt *RAB13* 3' UTR F | GGAATCTAGAGGACCCTTTCTTGCCTCCCCA |
| XbaI 90nt *RAB13* 3' UTR F | GGAATCTAGATGGAGGGTCACATAGGTA |

| | |
|---|---|
| XbaI 196nt *RAB13* 3' UTR F | GGAATCTAGATGAATTGAGGAAGTGAAAGAAGGC |
| XbaI 283nt *RAB13* 3' UTR F | GGAATCTAGATGGGTTTTCAGGGCAAAC |
| XhoI 89nt *RAB13* 3' UTR R | GGAACTCGAGAGCCCCTCTGCTATTTCTCC |
| XhoI 195nt *RAB13* 3' UTR R | GGAACTCGAGTTCCTCTCCCTTCTCCCTTC |
| XhoI 282nt *RAB13* 3' UTR R | GGAACTCGAGGGTAAGGTCTGAAGCCTGAG |
| XhoI 482nt *RAB13* 3' UTR R | GGAACTCGAGCTGACAATAATCCAAGTTACCC |
| del90-282nt *RAB13* 3' UTR F | GGGAGAAATAGCAGAGGGGCTTGGGTTTTCAGG |
| del90-282nt *RAB13* 3' UTR R | CTGAAAACCCAAGCCCCTCTGCTATTTCTCCC |
| del89-138nt *RAB13* 3' UTR F | GAGAAATAGCAGAGGGGCAAAGGGAAAAGCAGAAAG |
| del89-138nt *RAB13* 3' UTR R | CTTTCTGCTTTTCCCTTTGCCCCTCTGCTATTTCTC |
| del138-185nt *RAB13* 3' UTR F | GAATGAGGAGAAAAAGGAGGGGAGAGGAATGAATTGAG |
| del138-185nt *RAB13* 3' UTR R | CTCAATTCATTCCTCTCCCCTCCTTTTTCTCCTCATTC |
| del186-233nt *RAB13* 3' UTR F | GAGAGAGGAAGGGAGAAAGAGGGAGGAGGAAA |
| del186-233nt *RAB13* 3' UTR R | TTTCCTCCTCCCTCTTTCTCCCTTCCTCTCTC |
| del234-282nt *RAB13* 3' UTR F | GCAAGGAGGTAGGAAGTGGGTTTTCAGGGCAA |
| del234-282nt *RAB13* 3' UTR R | TTGCCCTGAAAACCCACTTCCTACCTCCTTGC |

**Notes:** Bold lowercase - T7 promoter; underlined lower case - tail-annealing sequence; underlined uppercase - restriction site; lowercase - Gateway *att*B site.

### Extended Data Figure 1

**a**

| Cluster | GO Cellular Component | % | *P* value | GO Molecular Function | % | *P* value |
|---|---|---|---|---|---|---|
| *k* 1 | intracellular organelle lumen<br>organelle lumen<br>membrane-enclosed lumen | 63<br>63<br>63 | 2.66E-03<br>2.66E-03<br>3.08E-03 | structural constituent of ribosome<br>structural molecule activity | 44<br>44 | 1.35E-02<br>2.46E-02 |
| *k* 2 | protein complex | 20 | 4.72E-02 | enzyme binding<br>sequence-specific double-stranded DNA binding | 40<br>20 | 2.71E-03<br>2.86E-02 |
| *k* 3 | cytoskeletal part<br>cytoskeleton | 27<br>27 | 3.72E-02<br>4.44E-02 | phosphoric ester hydrolase activity<br>phosphatase activity<br>protein dimerization activity | 20<br>20<br>27 | 9.16E-03<br>9.16E-03<br>3.16E-02 |
| *k* 4 | N/A | | | N/A | | |
| *k* 5 | cell periphery<br>plasma membrane | 80<br>80 | 7.21E-03<br>7.21E-03 | enzyme binding | 60 | 4.83E-02 |
| *k* 6 | N/A | | | transcription factor activity, sequence-specific DNA binding<br>nucleic acid binding transcription factor activity<br>RNA polymerase II tf activity, sequence-specific DNA binding | 29<br>29<br>21 | 1.95E-02<br>1.95E-02<br>2.86E-02 |
| *k* 7 | proteinaceous extracellular matrix | 67 | 2.67E-02 | receptor binding<br>heparin binding<br>sulfur compound binding | 100<br>67<br>67 | 2.63E-03<br>1.95E-02<br>1.95E-02 |
| *k* 8 | cytosolic part<br>cytosolic ribosome<br>ribosomal subunit | 82<br>82<br>82 | 3.28E-17<br>3.28E-17<br>1.65E-16 | RNA binding<br>structural constituent of ribosome<br>structural molecule activity | 87<br>79<br>79 | 6.32E-15<br>1.53E-14<br>2.46E-13 |

**b**

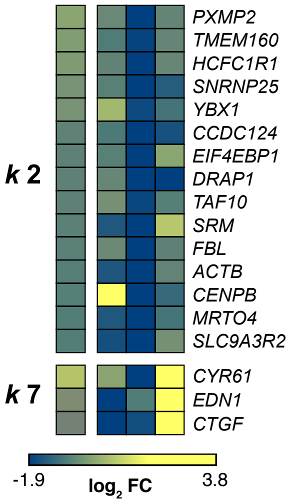


**c**

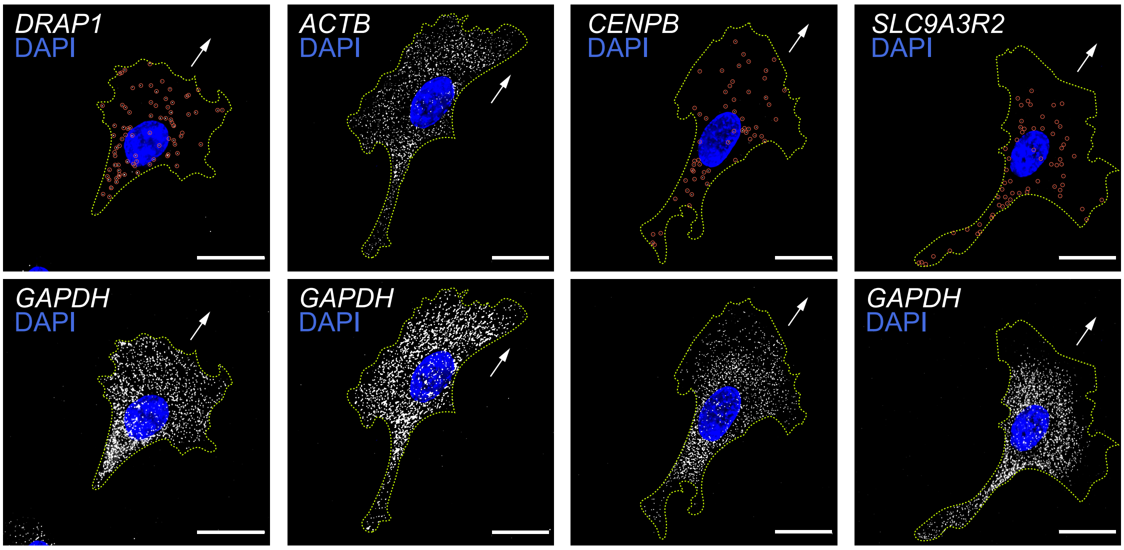


**k 2**

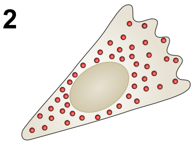

**d**

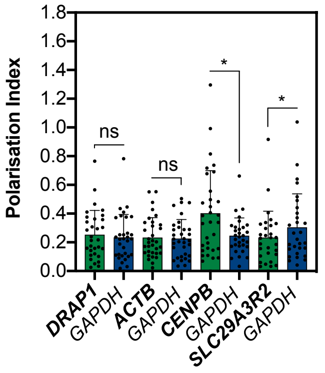


**e**

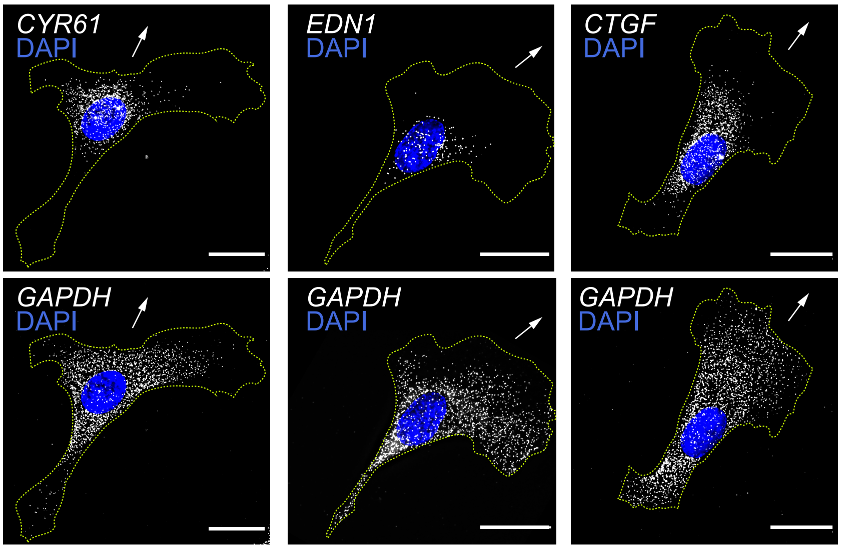


**k 7**

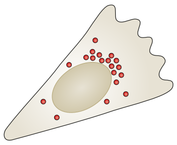

**f**

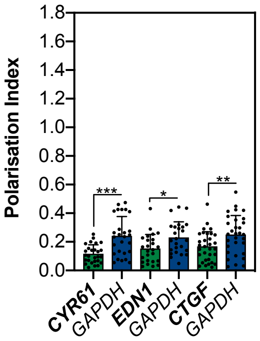

### Extended Data Figure 2

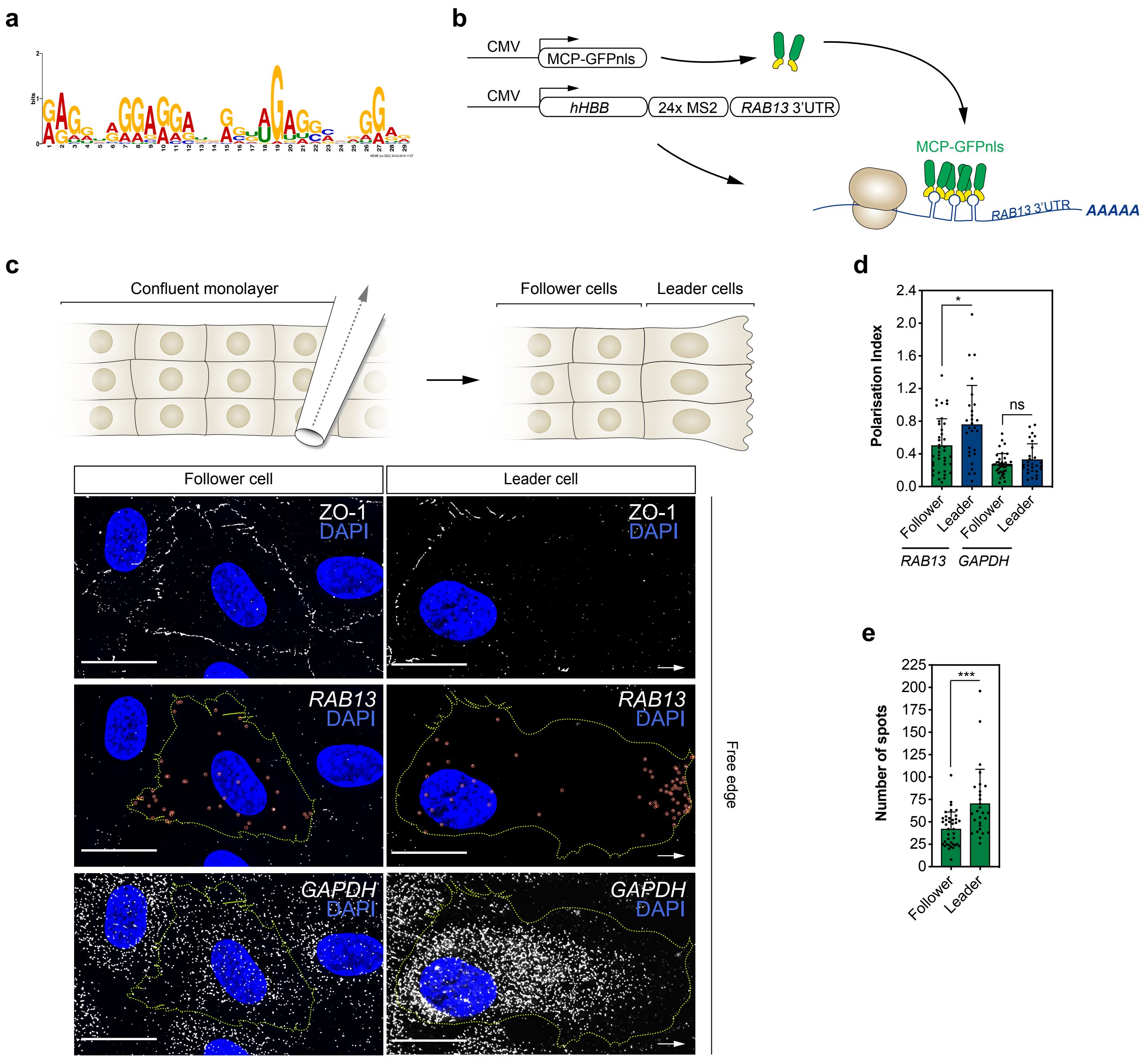

### Extended Data Figure 3

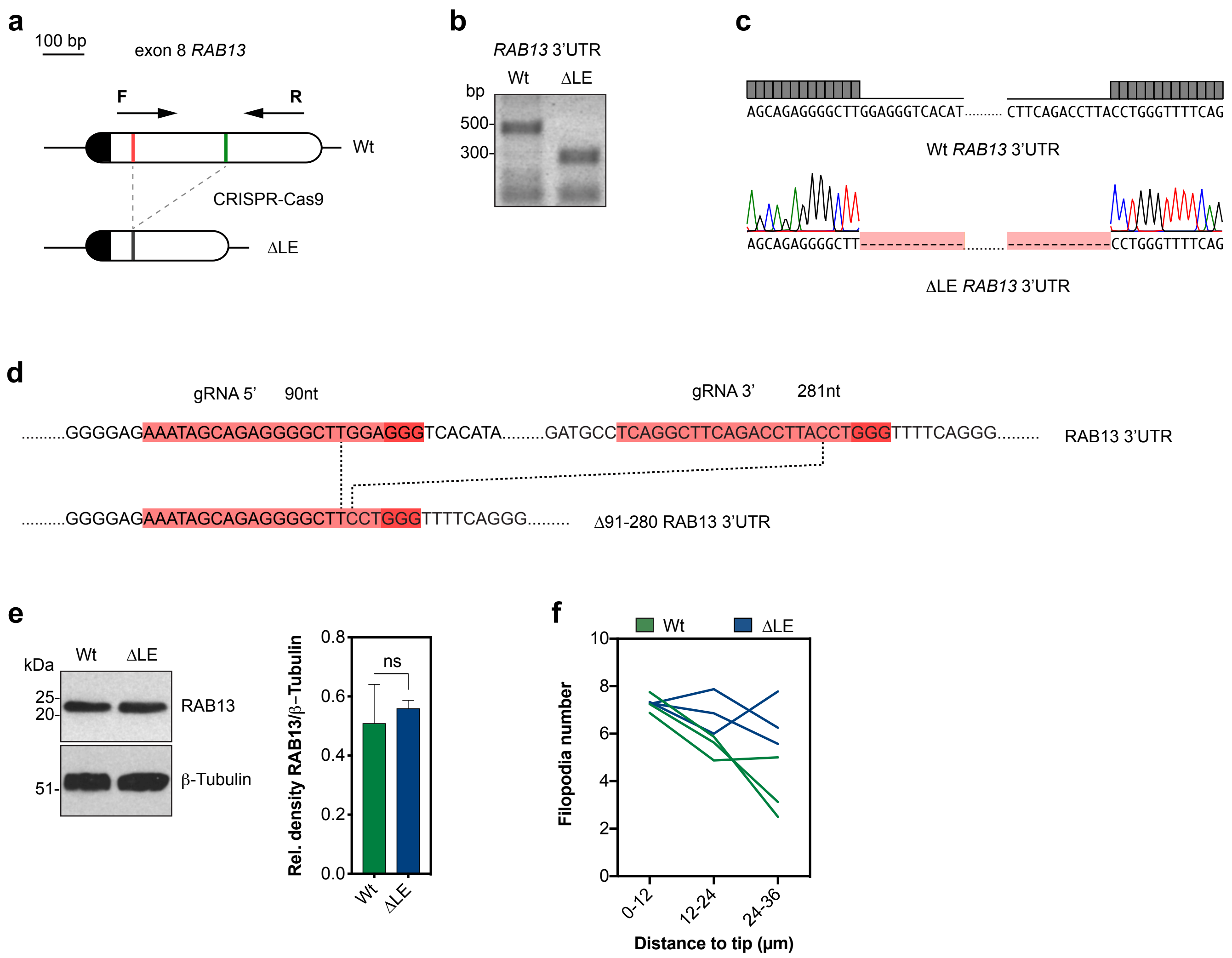

#### Extended Data Figure 4

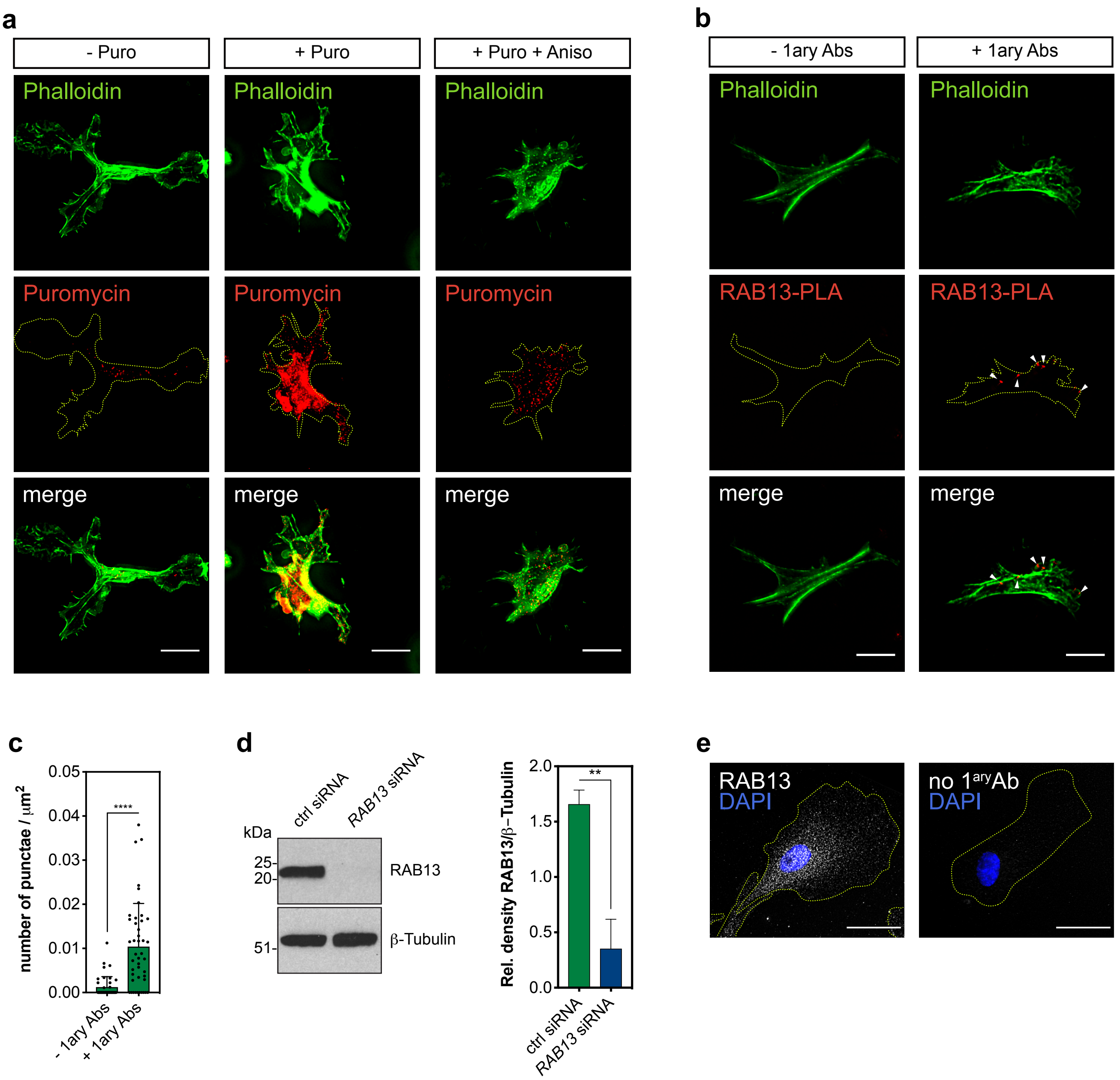

### Extended Data Figure 5

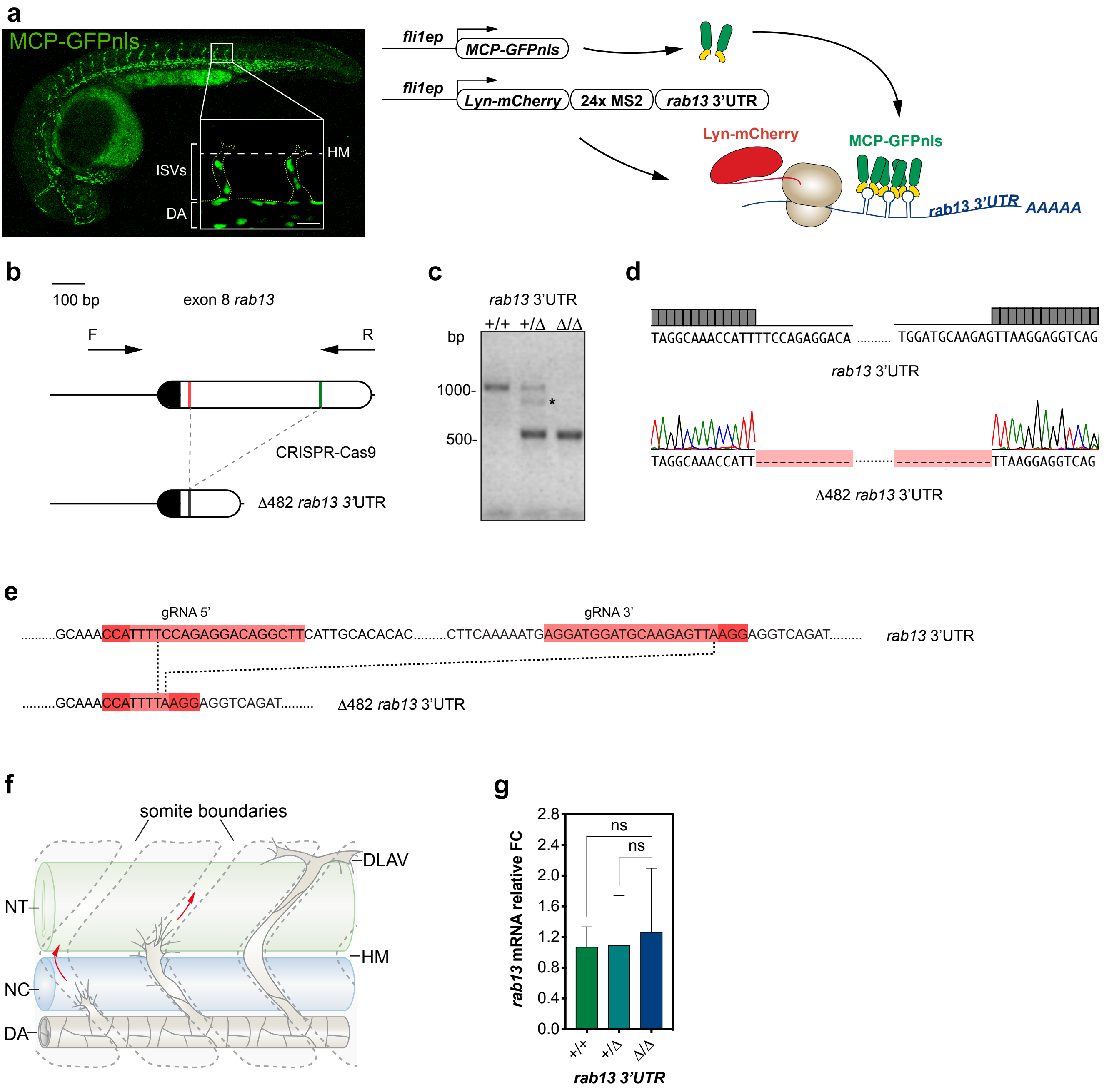

**SUPPLEMENTARY INFORMATION**

**Supplementary Video 1.** Confocal time-lapse imaging of a representative endothelial cell co-transfected with plasmids expressing Lyn-mCherry, MCP-GFPnls and 24xMS2-*RAB13* 3'UTR. Arrowheads represent newly-generated filopodia.

**Supplementary Video 2.** Time-lapse confocal micrographs of an ISV sprouting cell in a *Tg(fli1ep:MCP-GFPnls)* zebrafish embryo showing the localisation of MS2 hairpin-tagged *rab13* 3'UTR. Arrowheads indicate non-nuclear localisation of MCP-GFPnls.
